# Distal CA1 maintains a more coherent spatial representation than proximal CA1 when local and global cues conflict

**DOI:** 10.1101/2020.10.17.343558

**Authors:** Sachin S. Deshmukh

## Abstract

Entorhinal cortical projections show segregation along the transverse axis of CA1, with the medial entorhinal cortex (MEC) sending denser projections to proximal CA1 (pCA1) and the lateral entorhinal cortex (LEC) sending denser projections to distal CA1 (dCA1). Previous studies have reported functional segregation along the transverse axis of CA1 correlated with the functional differences in MEC and LEC. pCA1 shows higher spatial selectivity than dCA1 in these studies. We employ a double rotation paradigm, which creates an explicit conflict between local and global cues, to understand differential contributions of these reference frames to the spatial code in pCA1 and dCA1. We show that pCA1 and dCA1 respond differently to this local-global cue conflict. pCA1 shows incoherent response consistent with the strong conflicting inputs it receives from MEC and distal CA3 (dCA3). In contrast, dCA1 shows a more coherent rotation with global cues. In addition, pCA1 and dCA1 display comparable levels of spatial selectivity in this study. This finding differs from the previous studies, perhaps due to richer sensory information available in our behavior arena. Together these observations indicate that the functional segregation along proximodistal axis of CA1 is not merely of the amount of spatial selectivity but that of the nature of the different inputs utilized to create and anchor spatial representations.

## Introduction

The hippocampus is involved in spatial navigation and episodic memory (O’keefe and Nadel 1978; Squire et al. 2004). To understand the computations involved in these processes, it is critical to understand information transformation in the sub-regions of the entorhinal-hippocampal network. Cortical information to the hippocampus gets channeled through the medial and the lateral entorhinal cortex (MEC and LEC) (Burwell 2000; Witter and Amaral 2004). MEC and LEC are thought to convey complementary information to the hippocampus - path integration derived spatial information from MEC and nonspatial information from LEC (Deshmukh and Knierim 2011; Hafting et al. 2005; Knierim et al. 2014; Manns and Eichenbaum 2009; Suzuki et al. 1997); recent studies have demonstrated object dependent allocentric (Deshmukh and Knierim 2011) and egocentric (Wang et al. 2018) representations of space in LEC. While LEC and MEC layer II inputs to the dentate gyrus and CA3 are not strongly segregated along the transverse axis of the hippocampus, layer III inputs to CA1 are. MEC projects preferentially to proximal CA1 (pCA1; close to CA2), while LEC projects preferentially to distal CA1 (dCA1; close to the subiculum) (Naber et al. 2001; Steward and Scoville 1976; Witter and Amaral 2004). CA3 to CA1 projections also show segregation along the transverse axis, with proximal CA3 (pCA3) projecting preferentially to dCA1 and distal CA3 (dCA3) projecting preferentially to pCA1 (Figure 1A) (Ishizuka et al. 1990; Witter and Amaral 2004).

**Figure 1.**
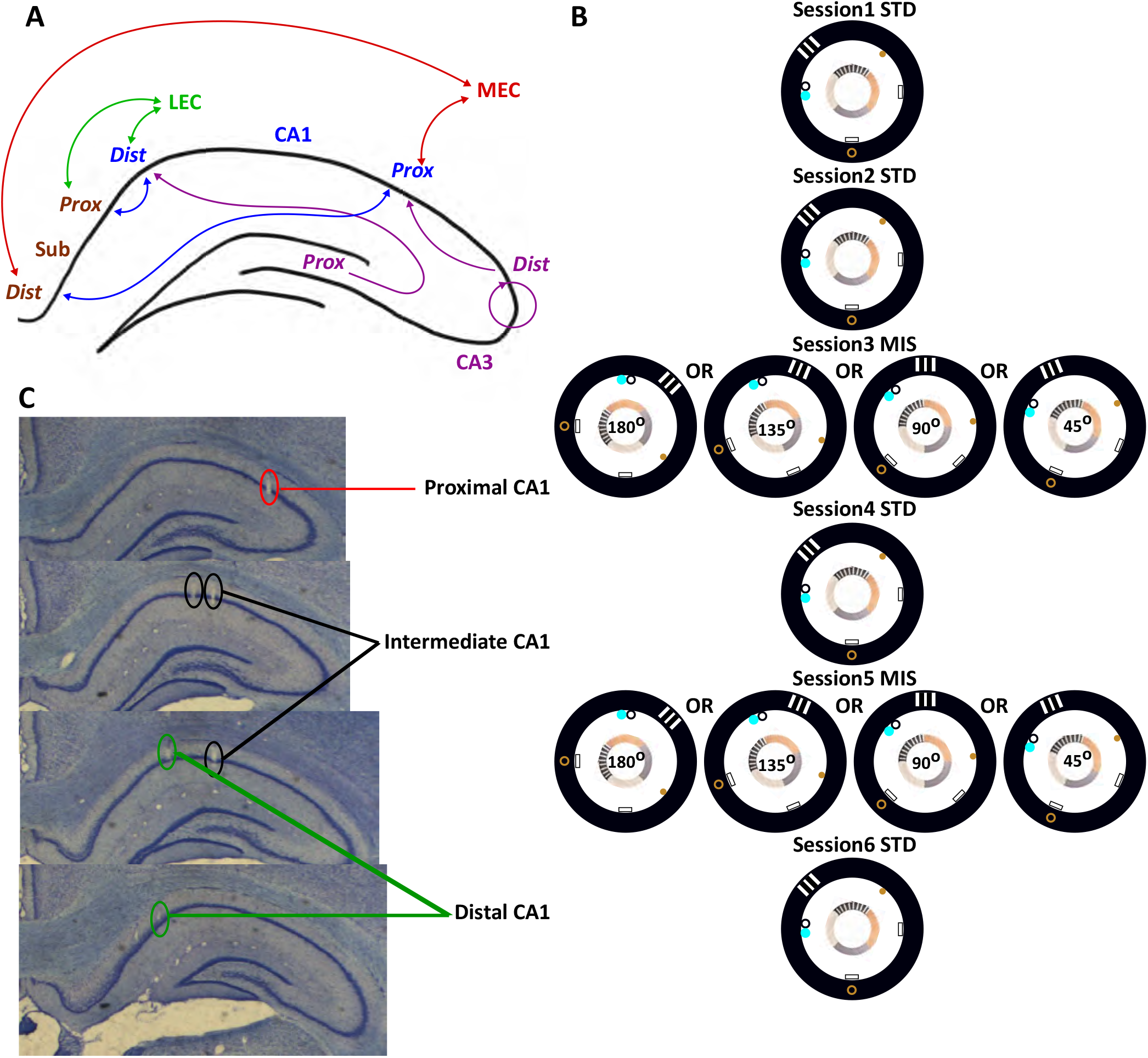
CA1 circuitry and experimental paradigm. (A) Schematic showing anatomical connectivity of CA1 along its transverse axis. Bidirectional arrows indicate reciprocal connections between the two connected areas; unidirectional arrows indicate direction of information flow. (B) Recording sessions included two mismatch sessions interleaved between standard sessions. All but one rat encountered 2 standard sessions before manipulations began for a total of 6 sessions each day, while one rat encountered only one standard session before manipulations began for a total of 5 sessions each day. Standard sessions had local as well as global cues in the configuration the rats were trained on, while mismatch sessions had local cues rotated CCW and global cues rotated CW by equal amounts to get a net cue mismatch of 45°, 90°, 135°, or 180°. (C) Examples of tetrodes recording along the transverse axis of CA1 from one rat. The drive canulae were linearly organized in ~ 3 rows oriented at 35° to the ML axis to target the entire extent of the transverse axis at the same septotemporal level.

As predicted by these anatomical differences, CA1 shows functional segregation along its transverse axis. In two-dimensional open fields and linear tracks, pCA1 is spatially more selective than dCA1 (Henriksen et al. 2010; Ng et al. 2018; Oliva et al. 2016). Consistent with the predictions, dCA1 neurons respond to objects and rewards (Burke et al. 2011; Xiao et al. 2020; these studies did not record from pCA1, so we do not know if there was a functional dissociation along the transverse axis). Social place cells, encoding location of a conspecific are more prevalent in dCA1 than pCA1 of bat (Omer et al. 2018). LEC and dCA1 show enhanced oscillatory synchronization during olfactory-spatial associative memory, while LEC and pCA1 do not (Igarashi et al. 2014). Immediate early gene expression and lesion studies lend further support to spatial vs non spatial double dissociation between pCA1 and dCA1 (Ito and Schuman 2012; Nakamura et al. 2013; Nakazawa et al. 2016). However, the relative contributions of different inputs to the neural representations along the transverse axis of CA1 are not well understood.

A “double rotation” protocol (Knierim 2002; Shapiro et al. 1997) has been used extensively to study the influence of local and global cues on spatial representations in different parts of the hippocampal formation. In this experimental protocol, prominent “local” on-track cues and “global” off-track cues are put in conflict by rotating them in opposite directions. These experiments have revealed that CA1 place fields respond to double rotation in an incoherent manner, while CA3 place fields rotate coherently with the local cues (Lee et al. 2004; Neunuebel and Knierim 2014). Along the transverse axis of CA3, pCA3 place cells show an incoherent response while intermediate CA3 (iCA3) and dCA3 place cells show coherent rotation with the local cues (Lee et al. 2015). In the same paradigm, MEC shows coherent rotation with the global cues. LEC, which shows very weak spatial tuning during the sessions with the standard cue configuration, shows weak (but statistically significant) rotation with the local cues (Neunuebel et al. 2013). Thus, pCA1 gets strongly coherent but conflicting inputs from MEC (global) and dCA3 (local), while dCA1 gets incoherent inputs from pCA3 and weakly local inputs from LEC. The present study examines responses of CA1 neurons along the transverse axis to the double rotation manipulation to study how the entorhinal and CA3 input patterns affect CA1 representation along its transverse axis during local-global cue conflict. Our results show that pCA1, which receives strong but conflicting inputs from MEC and dCA3, shows lower coherence in its response to double rotation compared to dCA1. Furthermore, spatial selectivity in pCA1 and dCA1 is comparable in this experimental paradigm. Thus, dCA1 is not necessarily less spatial than pCA1, as claimed in the earlier reports (Henriksen et al. 2010; Ng et al. 2018; Oliva et al. 2016).

## Methods

### Subjects and surgery

Eleven Long Evans rats aged 5-6 months were housed individually on a 12:12 hour reversed day-night cycle. All experiments were performed during the night portion of the cycle. Animal care, surgical and euthanasia procedures were in accordance with the National Institutes of Health guidelines and protocols approved by the Institutional Animal Care and Use Committee of the Johns Hopkins University. Custom built hyperdrives with independently moving 15 tetrodes and 2 references were implanted over the right hemisphere. Rats were implanted with 3D printed hyperdrives with linearly distributed bundle canulae angled at 35° to the ML axis to target the entire proximodistal extent of CA1 at the same septotemporal level. Data recorded during resting sessions from 8 of these rats was previously used for studying propagation of ripples in CA1 (Kumar and Deshmukh 2020).

### Behavioral training and experimental protocol

Following recovery for about one week after surgery, rats were maintained at 80-90% of their free feeding weight to incentivize them to run on a circular track for food, during the training and the recording sessions. The circular track (56 cm inner diameter, 76 cm outer diameter) had 4 easily distinguishable sections, and was placed in a room with a 2.75 m diameter circular curtain with 6 large, distinct cues either hanging on the curtain or on the floor along the curtain (Knierim 2002). Once the rats learned to run in clockwise (CW) direction for food pellets randomly placed on the track, and the electrodes were deemed to be in optimal recording locations, the experimental sessions commenced. Each experimental day had 5-6 sessions of 15 laps each. The experimental sequence for one rat was STD-MIS-STD-MIS-STD, while that for all other rats was STD-STD-MIS-STD-MIS-STD. STD stands for standard configuration, and MIS stands for mismatch configuration with one out of the following mismatch angles selected in a pseudorandom order: 45°, 90°, 135°, and 180°. For a given mismatch angle, the local cues on the track were rotated counterclockwise (CCW) by half the amount and the global cues along the curtain were rotated CW by the other half. For example, the local cues were rotated by 22.5° CCW and the global cues were rotated by 22.5° CW for a 45° MIS session (Figure 1B). These manipulations were performed for 4 days, such that each mismatch angle was sampled twice.

### Recording electronics

Neuronal data was collected using an analog wireless transmitter (Triangle Biosystems International, Durham, NC). Data from the wireless receiver was processed and stored using cheetah data acquisition system (Neuralynx Inc., Bozeman, MT). The signals were amplified 1000-10000 fold, bandpass filtered between 600-6000 Hz, and digitized at 32000 Hz for single unit recordings. Any time one of the channels on a tetrode crossed a preset threshold, data from all 4 channels on the tetrode were recorded for 1ms (8 samples before and 24 samples after the threshold). Signals from one of the channels on each of the tetrodes were also amplified 500-2000 fold, bandpass filtered between 1-475 Hz, digitized at 1 KHz and stored continuously for local field potential (LFP) recordings.

### Data analysis

#### Unit isolation

Single units were isolated using WinClust, a custom manual cluster cutting software (J. J. Knierim, Johns Hopkins University). For every threshold crossing, waveform characteristics such as peak, valley and energy on all 4 channels on a tetrode were used for clustering spikes. Only units with fair or better isolation as estimated by cluster separation, waveform and clean inter-spike interval histogram were included in subsequent analysis. Putative interneurons firing at ≥ 10 Hz mean firing rate were excluded.

#### Firing rate maps

Rat’s position as well as heading direction were tracked using colored LEDs and a camera recording at 30 frames per second. Off track firing and intervals during which the rats ran slower than 2 cm/s or ran in the wrong direction (CCW) were excluded from the analysis to minimize firing rate variability introduced by non-spatial activity. Linearized rate maps were created at 1° resolution (which gives 360 bins for the circular track) by dividing the number of spikes when the rat was in each bin by the amount of time the rat spent in each bin. Linearized rate maps were smoothed using adaptive binning for computing spatial information scores (bits/spike; Skaggs et al. 1996) and gaussian filtered (σ = 3°) for other quantitative analyses. A shuffling procedure was used to determine the probability of obtaining the observed spatial information score by chance. The neuronal spike train was shifted by a random lag (minimum 30 s) with respect to the trajectory of the rat. Spatial information score was computed from the adaptive binned, linearized rate map created from this shifted spike train. This procedure was repeated 1000 times to estimate the chance distribution of spatial information scores, and determine the number of randomly time shifted trials having spatial information scores greater than or equal to the observed spatial information scores. Only putative place cells that fired ≥ 20 spikes, and had statistically significant (p < 0.01) spatial information scores of ≥ 0.5 bits/spike in at least one of the MIS and preceding STD session were used in population vector correlation (PVC) analysis and single unit responses.

Peak firing rate was the maximum firing rate in the gaussian smoothed linearized rate map while the mean firing rate was mean of the firing rates in all bins of the same. For the neurons satisfying spatial information criteria and peak firing rates above 1 Hz, place fields were defined as regions with minimum 10 contiguous pixels with firing rates above 15% of the peak firing rate or 0.3 Hz, whichever was higher. Number of place fields meeting these criteria in each rate map were counted. Field size was the number of 1° pixels in each place field while fraction of track occupied by the place fields was the total number of 1° pixels occupied by all the place fields of a neuron normalized by 360 (total number of pixels in the rate map).

#### Classification of single unit responses

The angle of rotation giving the highest Pearson correlation between the STD and MIS sessions was used as an estimate of rotation of the neuron’s place field. We categorized responses of putative place cells as “appear” (< 20 spikes in STD but > 20 spikes in MIS), “disappear” (< 20 spikes in MIS but > 20 spikes in STD), “local” (highest Pearson correlation between STD and MIS after CCW rotation), “global” (highest Pearson correlation between STD and MIS after CW rotation), or “ambiguous” (Pearson correlation coefficient between STD and MIS not crossing a threshold of 0.6 after rotation, or the neuron not meeting the spatial information criteria in one of the sessions).

#### PVC analysis

Linearized rate maps were normalized by their peak firing rate before being used for constructing population vectors. Normalized firing rates of all cells for each 1° bin in each MIS or STD session constituted the population vector for that bin for that session. STD vs MIS PVC matrices were constructed by computing Pearson correlation coefficients of the 360 population vectors of each STD and MIS session with each other at all possible relative displacements (0-359°). Similarly, STD vs STD PVC matrices were constructed from STD sessions before and after MIS (Lee et al. 2004). The mean PVC at each relative displacement (0-359°) was computed from each PVC matrix. Peaks (polar (length, angle) pairs converted to cartesian (x, y) pairs), as well as full width at half maximum (FWHM) were estimated from the mean PVC. Normalized bias was defined as the difference between the mean PVCs for all displacements in the direction of local cue rotation (1-179°) and mean PVCs for all displacements in the direction of global cue rotation (181-359°) normalized by the mean PVC for all displacements (0-359°).

#### Shuffle analysis of PVCs

If a given STD vs MIS comparison had n pCA1 and m dCA1 neurons, these n + m neurons were randomly reassigned to shuffled pCA1 and dCA1 groups, with n and m neurons, respectively. Peak PVCs were obtained from the shuffled data as described above. This shuffling procedure was repeated 1000 times to create a peak PVC distribution of the shuffled data.

#### Bootstrap analysis of PVCs

We performed bootstrap analysis by random sampling with replacement of neurons in the dataset to generate 1000 resampled datasets with the number of samples matching the number of neurons in the actual dataset for each region. Peaks of mean PVCs (x,y), FWHM, and normalized bias were calculated for each of the bootstrap iterations. Because these different parameters have different units and different magnitudes, they were normalized by subtracting their minima and dividing by the difference between their maxima and their minima. This led to all parameters having a range of 0 to 1. K-means clustering was used to partition the bootstrap distributions from pCA1 and dCA1 into 2 clusters. Principal component analysis (PCA) was used to reduce dimensionality for display purpose.

#### Statistical analysis

MATLAB (Mathworks, Natick, MA) was used to perform statistical analysis. The circular statistics toolbox was used for statistical analysis of circular data. Units were tracked through a recording day, which typically had two MIS sessions with different mismatch angles. Tetrodes with multiple units were left undisturbed from one day to another, while those without units were moved ~16-32 μm in an attempt to increase the yield. For tetrodes with units on multiple days, no attempt was made to track units from one recording day to another. Therefore, while a number of units could be shared between sessions with different mismatch angles, we do not know the exact number of shared units. This partial overlap in number of units violates the assumption of independence made while correcting for multiple comparisons, such as Bonferroni or Holm-Bonferroni correction(Holm, 1979). Thus, rather than using corrections for multiple comparisons, patterns of low p values (p < 0.05) across multiple tests were used to draw conclusions. No conclusions were drawn based on single comparisons where multiple comparisons were performed simultaneously.

#### Histology

On the final day of recording, locations of a small subset of tetrodes were marked by passing 10 μA current for 10 s. Tetrode tracks were reconstructed from coronal sections and confirmed by the marker lesions (Figure 1C; Deshmukh et al. 2010).

## Results

Hyperdrives with 15 tetrodes and 2 references targeting the entire proximodistal extent of dorsal CA1 were implanted on 11 rats to record the activity of putative pyramidal cells as the rats ran CW on a circular track with 4 distinct textures (local cues) in a circular curtained room with 6 large global cues along the curtain. Cue manipulation sessions with global cues moving CW and local cues moving CCW by equal amounts creating 45°, 90°, 135°, or 180° mismatch (MIS) between the two were interleaved with sessions with cues in their standard configuration (STD) (Figure 1B).

Putative interneurons with mean firing rates > 10 Hz (Deshmukh and Knierim 2013; Fox and Ranck 1981; Frank et al. 2001; Ranck 1973) were excluded from analyses. Based on their locations, tetrodes were assigned to equally broad pCA1, intermediate CA1 (iCA1) and dCA1 bands (Henriksen et al. 2010). Nine rats had putative pyramidal neurons recorded from in each of the three bands; one rat had units recorded from pCA1 and iCA1; one rat had units recorded from pCA1 and dCA1. Since iCA1 is expected to have overlapping entorhinal projections (Naber et al. 2001; Steward and Scoville 1976), although data from all three regions is displayed, quantitative statistical comparisons between regions were limited to pCA1 and dCA1 by prior design.

### Properties of single units along the proximodistal axis of CA1

In the first STD session, 156 well isolated putative pyramidal cells in pCA1 fired at least 20 spikes while the rat was running on the track; 131 neurons in iCA1 and 180 neurons in dCA1 met the same criteria. Mean and peak firing rates in pCA1 and dCA1 were statistically indistinguishable from each other (Wilcoxon rank sum test, mean firing rate: pCA1 median = 1.24 Hz, dCA1 median = 1.29 Hz, p = 0.37; peak firing rate: pCA1 median = 11.97 Hz, dCA1 median = 11.52 Hz, p = 0.34). Spatial correlates of neural activity were estimated using a variety of measures. Spatial information scores (Skaggs et al. 1996) in pCA1 and dCA1 neurons were statistically indistinguishable from one another (pCA1 median = 1.92 bits/spike, dCA1 median = 2.11 bits/spike; Wilcoxon rank sum test, p = 0.15). Number of place fields/cell of 139 pCA1 and 140 dCA1 place cells with statistically significant spatial information scores > 0.5 bits/spike were similarly indistinguishable (median = 1 for both, Wilcoxon rank sum test, p = 0.28), and so were the fraction of place cells with a single place field (χ2 test, p = 0.36). Furthermore, sizes of 181 pCA1 and 176 dCA1 place fields were statistically indistinguishable (pCA1 median = 44°, dCA1 median = 47°; Wilcoxon rank sum test, p = 0.13), and so was the fraction of track occupied by all the place fields of pCA1 and dCA1 place cells(pCA1 median = 0.17, dCA1 median = 0.17; Wilcoxon rank sum test, p = 0.13). This lack of difference in spatial correlates of pCA1 and dCA1 (Figure 2; Supplementary Table 1 shows the detailed statistics for all single unit properties) persisted in MIS sessions irrespective of MIS angle. Spatial correlates showed a general decline from STD to MIS session in both regions (Supplementary Figure 1, Supplementary Table 2).

**Figure 2.**
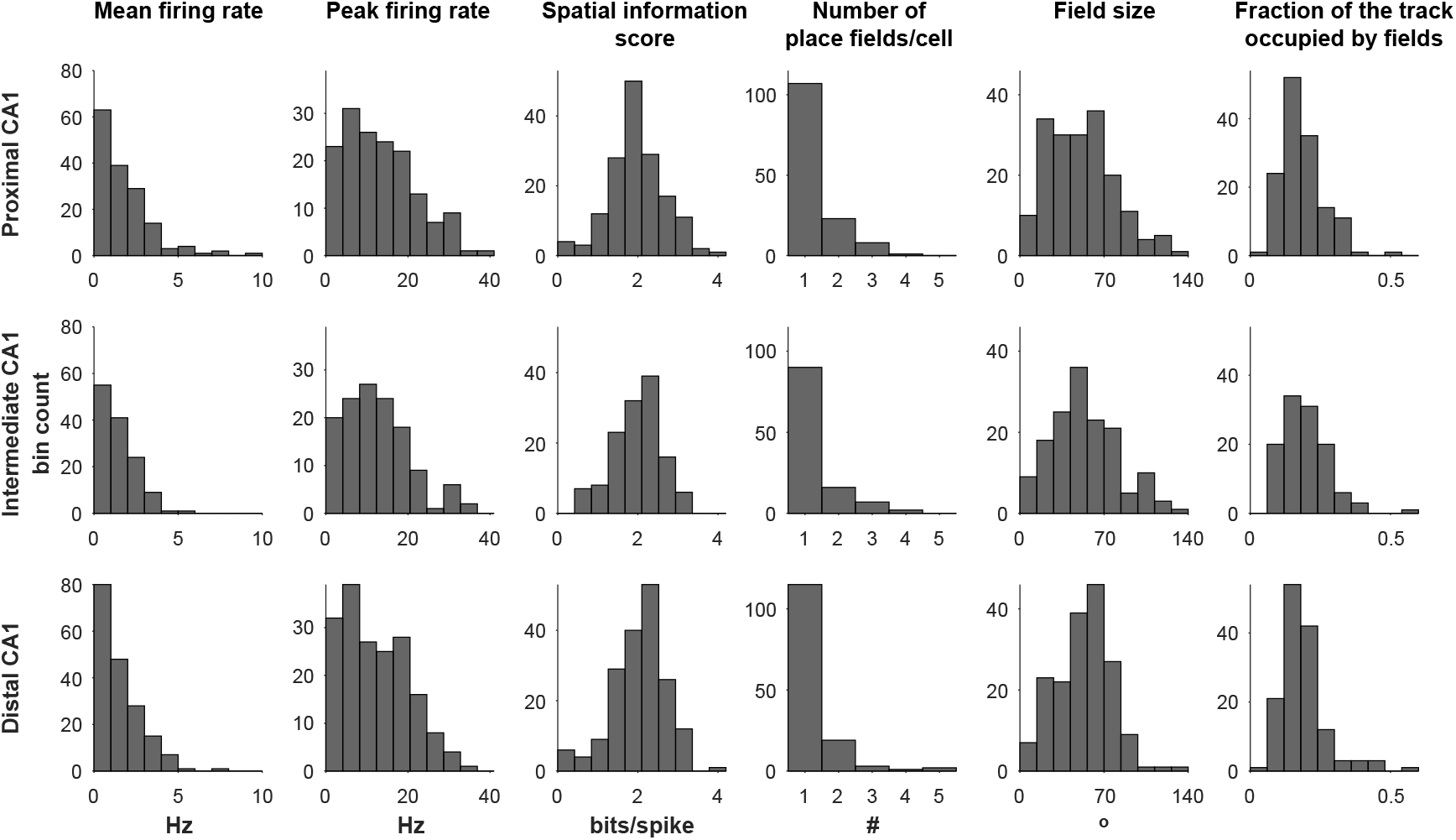
Properties of putative pyramidal cells along the transverse axis of CA1. Mean firing rates, peak firing rates, spatial information scores, number of place fields per cell, place field sizes, and fraction of the track occupied by place fields in pCA1, iCA1 and dCA1 during the first standard session of the day do not differ from one another.

### Single unit responses to cue manipulation

Responses to cue manipulation of putative place cells (≥ 20 spikes and spatial information score > 0.5 bits/spike in at least one of the STD or MIS session) were grouped into 5 classes as described previously from animals performing the same behavioral task (Figure 3A; Lee et al. 2004; Neunuebel et al. 2013). Units that remained spatially selective in both STD and MIS sessions and showed maximum Pearson correlation after rotating the rate maps > 0.6 were classified as rotating neurons – those rotating CW were classified as rotating with global cues, while those rotating CCW were classified as rotating with local cues. Units that dropped below the 20 spikes threshold in MIS session were classified as disappearing units, while those that started firing more than 20 spikes in MIS after firing less than that in STD were classified as appearing units. Units that failed to meet the maximum Pearson correlation coefficient criterion as well as those which failed the spatial information score criteria in at least one of the sessions while firing more than 20 spikes in both sessions were classified as being ambiguous. After pooling across all mismatch angles, proportions of appearing, disappearing, and ambiguous units were similar in pCA1 and dCA1, but pCA1 showed more units with CCW rotations in the direction of local cues while dCA1 showed more units with CW rotations in the direction of global cues (Figure 3B). Many neurons are likely to have been recorded in more than one mismatch angles. In addition, classification of neurons into response classes creates arbitrary distinctions. For these two reasons, we did not perform quantitative statistical analysis on the distributions of neurons in these classes. Nonetheless, this classification provides a useful qualitative description of single unit responses that underlie the quantitative differences in populations of neurons across the transverse axis of CA1 discussed below.

**Figure 3.**
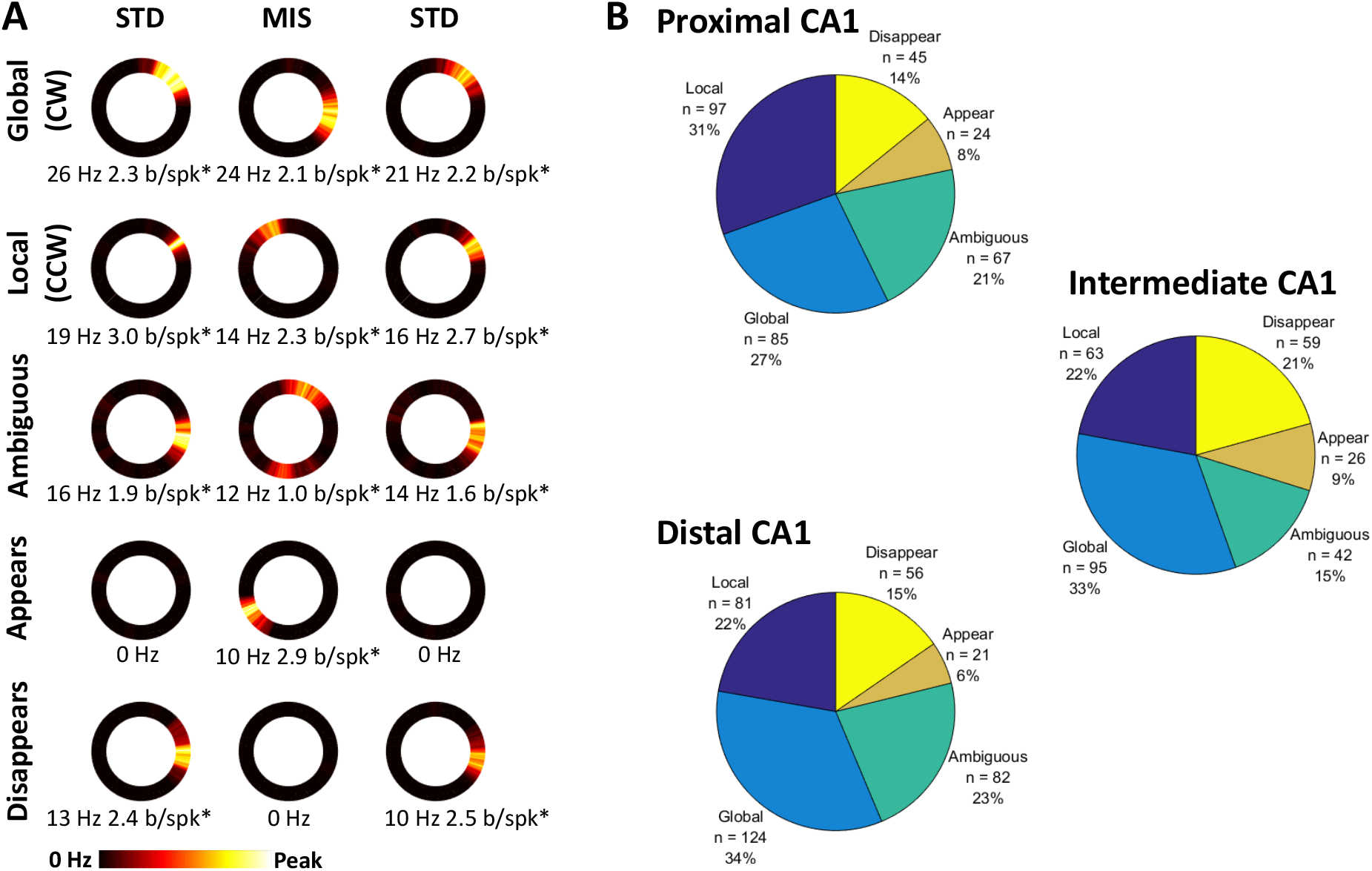
Single unit responses to cue manipulation. (A) Examples of units showing different types of responses to cue manipulations. Responses were categorized into 5 types: CW rotation in the direction of the global cues, CCW rotation in the direction of the local cues, ambiguous rotation, appear in MIS session and disappear in MIS session. Peak firing rates and spatial information scores are shown under each rate map. Asterisks mark statistically significant spatial information scores. (B) Proportion of different classes of responses in pCA1, iCA1 and dCA1 pooled across the 4 mismatch angles.

Rotating cells showed clustering of rotation angles near the rotation angles of local or global cues in MIS sessions (Figure 4). These rotation angles were not distributed uniformly in 0-360° range for any of the mismatch angles for any of the CA1 subregions (Rao’s spacing test, p < 0.001 for all angles for all regions; Supplementary Table 3), confirming that they were not distributed by a random process with uniform distribution. Coherence of rotation of the rotating cells was estimated using mean vectors (Rayleigh test, Supplementary Table 4). pCA1 single units showed statistically significant coherent rotations at all mismatch angles. While the mean vectors (blue arrows in figure 4A) were rotated CCW towards local cues for 90°, 135° and 180°, the mean vector for 45° was rotated CW towards global cues. In contrast, mean vectors of dCA1 single units showed CW rotations towards global cues for all mismatch angles. The mean vectors were statistically significant for all mismatch angles except 180°. Mean vector length and Rayleigh’s test used to measure its significance are flawed when the distribution is clearly bimodal, as seen in Figure 4A. A bimodal distribution gives shorter mean vector lengths, especially when the two modes are 180° apart, even when the distribution is clearly non-random (as confirmed by Rao’s spacing test). There was no discernible pattern in the angles of rotation of mean vectors of single units in iCA1, and the mean vectors were statistically significant only at 45°and 90°. Across all mismatch angles, pCA1 showed similar proportion of units rotating CW towards global cues (47%) and CCW towards local cues (53%). In contrast, both distal and iCA1 showed a preference for CW rotation towards global cues (60% vs 40% CCW) (Figure 4B).

**Figure 4.**
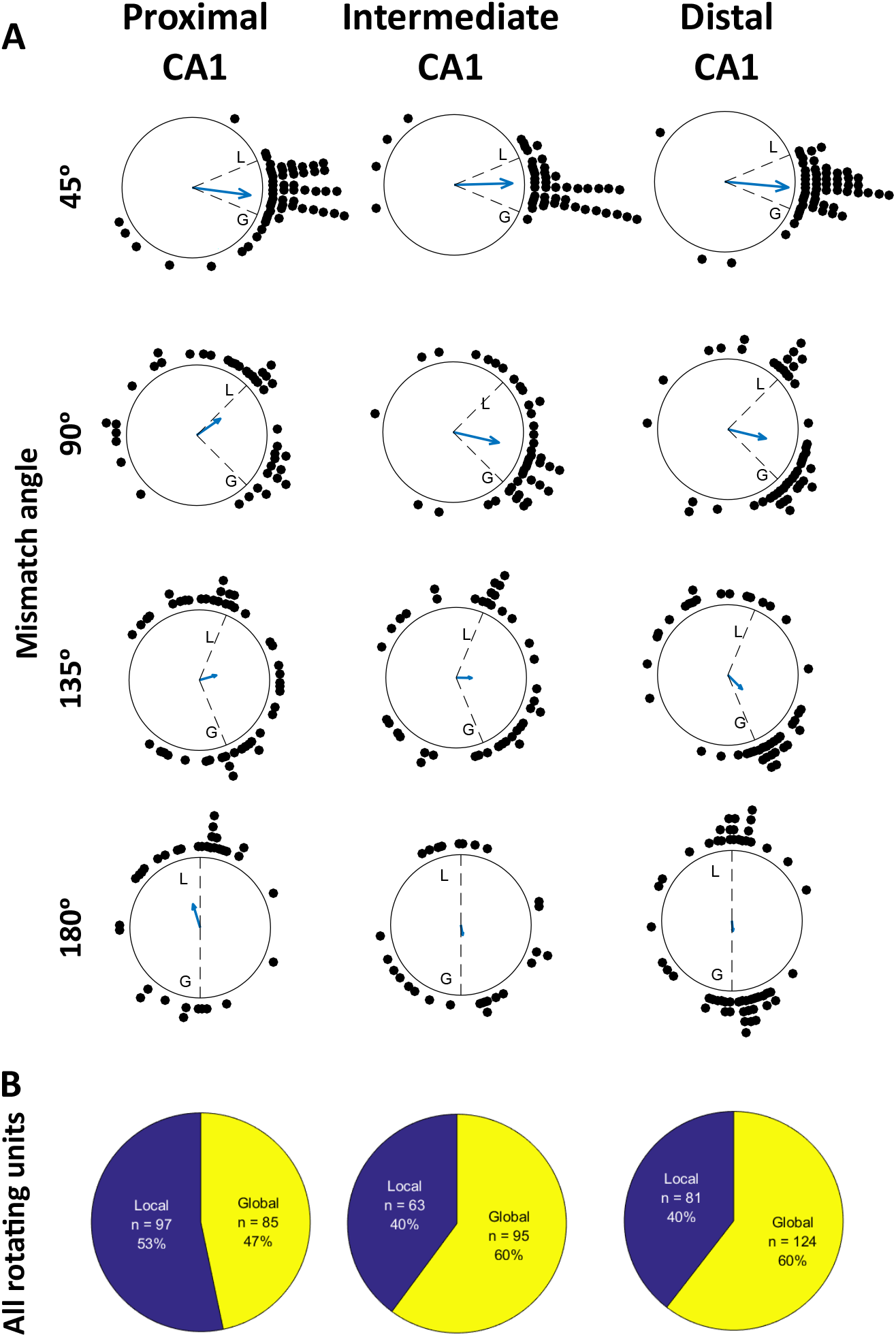
Distribution of single cell rotation angles in response to increasing mismatch angles. (A) Angle of rotation of each rotating unit between STD and MIS sessions is represented by a dot around a circle. Mean vector computed from the rotation angles of all units for the given MIS angle is represented by a blue arrow in the center. Angles of rotations of cues between STD and MIS sessions are represented by dotted lines; local and global cue rotations are marked by the letters L and G. (B) Pie charts showing proportions of rotating units summed across all mismatch angles.

### Population responses to cue manipulation

While the rotating single units in pCA1 and dCA1 demonstrate clear differences, single unit rotation analysis misses out on contributions of other neurons in the ensemble excluded by the criteria used for classifying single unit responses to cue manipulations as rotating. PVCs (Lee et al. 2015, Lee et al. 2004; Neunuebel et al. 2013) between sessions using all neurons meeting inclusion criteria (statistically significant spatial information scores > = 0.5 bits/spike, minimum 20 spikes, mean firing rate < 10 Hz) in either MIS or preceding STD session help overcome this limitation. PVCs between STD sessions preceding and following MIS sessions (labelled STD1 and STD2) were used as control.

Figure 5 shows PVCs for STD1 vs STD2 and STD1 vs MIS for all MIS angles for pCA1, iCA1 and dCA1. The strength of correlation between the population vectors as estimated by Pearson correlation coefficients at all combinations of relative displacements is represented in pseudocolor, with black corresponding to <= 0 and white corresponding to 1. If the population representation is unchanged between the two sessions being compared (STD1 vs STD2 or STD1 vs MIS), the population vectors are expected to show highest correlation at 0° displacement from one another. This generates a line of highest correlation at the diagonal going from bottom left to top right of the PVC matrix (black line). If, on the other hand, the population representation coherently rotates between the two sessions being compared, the line of highest correlation (running at 45° angle, parallel to the diagonal) in the PVC matrix is expected to get displaced by the corresponding angle. Colored lines in each PVC matrix show the expected displacement for the given MIS angle for local (red) and global (green) rotation. STD1 vs STD2 PVC matrices for all MIS angles and CA1 subregions show a strong band of high correlation at the diagonal overlapping with the black line. This indicates that the population representations along the entire extent of CA1 transverse axis remained stable between STD sessions before and after the intervening MIS session regardless of the mismatch angle. In the 45° MIS session, all three regions showed strong PVC bands, which were slightly displaced towards the green line corresponding to global cue rotation. In 90°, 135° and 180° MIS sessions, pCA1 did not show a single coherent band of high correlation parallel to the diagonal. Instead, patches of high correlations were seen overlapping with the local (red line) as well as global (green line) cue rotations. iCA1 showed a more coherent band of high correlation in the 90° MIS session, which was biased towards the global cue rotation (green line) but showed a patchy distribution of high correlations similar to pCA1 in 135° and 180° MIS. In contrast to these two regions, dCA1 showed a fairly coherent band of high correlation biased towards global cue rotation (green line) at all mismatch angles with occasional patches of high correlation near local cue rotations (red line).

**Figure 5.**
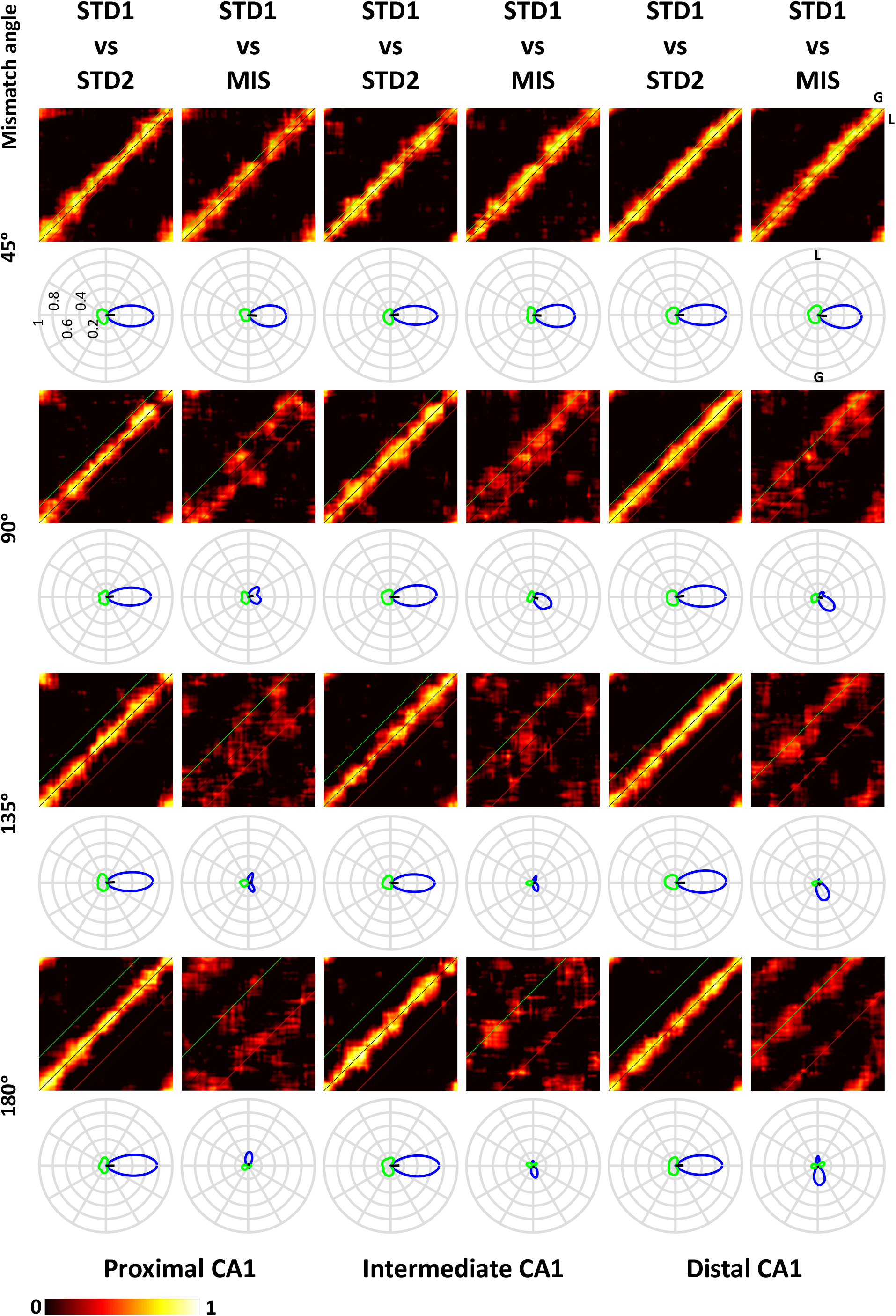
Population vector correlations. PVC matrices for STD1 vs STD2 sessions (STD sessions preceding and following MIS session) and STD1 vs MIS session show magnitude of correlation between population vectors for each 1° position bin on the track as a function of relative displacement between the two sessions being compared. STD1 vs STD2 matrices show a strong band of high correlation around 0° displacement (black line) for all mismatch angles and all subregions of CA1. Red and green lines show the expected displacement of high correlation band corresponding to local (L) and global (G) cue rotations in the different mismatch sessions, respectively. STD1 vs MIS matrices show different responses at different mismatch angles for different regions. While dCA1 STD1 vs MIS matrices show a distinct band of high correlation following global cue rotation (green line) at all mismatch angles, pCA1 and iCA1 STD1 vs MIS matrices show patchy distribution of high correlation at multiple mismatch angles. Polar plots below the PVC matrices show mean of PVCs over all positions at each relative displacement (0°-359°). Magnitudes of positive mean correlations are shown in blue while magnitudes of negative mean correlations are shown in green. Black line in the center shows mean vectors computed from the polar plots. Notice the reduced peak correlations at higher mismatch angles for all regions compared to STD1 vs STD2 correlations and STD1 vs MIS correlations at 45°. Contrast the bimodal structure of positive correlations for pCA1 at 90° and 135° with the larger peak following the global cues for dCA1. The outermost circle in the polar plot corresponds to Pearson correlation coefficient of 1 with each concentric circle being spaced at 0.2.

To quantify the strength of the correlation bands, the mean PVC for each relative displacement (0°-359°) was calculated from 360 correlation bins along a line running parallel to the bottom left to top right diagonal. Displacement of the line from the diagonal corresponded to the relative displacement between the two sessions (STD1 vs STD2 or STD1 vs MIS). Mean PVCs (plotted on polar plots below each PVC matrix in Figure 5) show a narrow distribution with large peaks at or near 0° displacement for STD1 vs STD2 comparisons for all CA1 subregions, as expected when the representations remain stable between sessions being compared. Peaks in the polar plots of all STD1 vs MIS comparisons were smaller than the corresponding STD1 vs STD2 comparisons for all CA1 subregions. For STD1 vs MIS comparison for the 45° mismatch angle, the distribution of mean PVCs widened and showed a bias towards CW rotations (in the direction of global cue rotation) in all CA1 subregions. As expected from the PVC matrices, the polar plot for higher mismatch angles showed different patterns in different subregions of CA1. pCA1 showed two comparable peaks of substantially reduced magnitude following local as well as global cue rotations in the 90° and 135° MIS sessions, and a prominent but small peak rotating CCW (following local cue rotation) in the 180° MIS session. iCA1 showed peaks rotating CW (following global cue rotation) in the 90° and 180° MIS sessions, and a two-peaked distribution in the 135° MIS session. dCA1, in contrast to the other two regions, showed consistent CW rotation with global cues in the 90°, 135° and 180° MIS sessions, although substantially smaller peaks corresponding to local cue rotations could also be seen.

We used a shuffle analysis to test if the neurons in pCA1 or dCA1 rotate more coherently than the combined population of the two regions. For each of the 1000 shuffles, neurons were randomly reassigned to pCA1 or dCA1 group while keeping the number of neurons in each group the same as the actual data. Peaks in the mean PVCs were determined for each group. The distribution of 1000 peak PVCs obtained for each of the groups simulates the distribution expected if the pCA1 and dCA1 neurons came from a common pool with comparable propensities to respond to the double rotation manipulation. A higher peak vector correlation than expected from this distribution indicates that the given region rotates more coherently than the common pool. Figure 6A shows that dCA1 consistently has higher correlation than 95^th^ percentile of the shuffled distribution across all MIS angles (45°: p = 0.019; 90°: p = 0.026; 135°: p = 0.012; 180°: p < 0.001) while pCA1 does not (45°: p = 0.953; 90°: p = 0.938; 135°: p = 0.983; 180°: p = 0.297).

**Figure 6.**
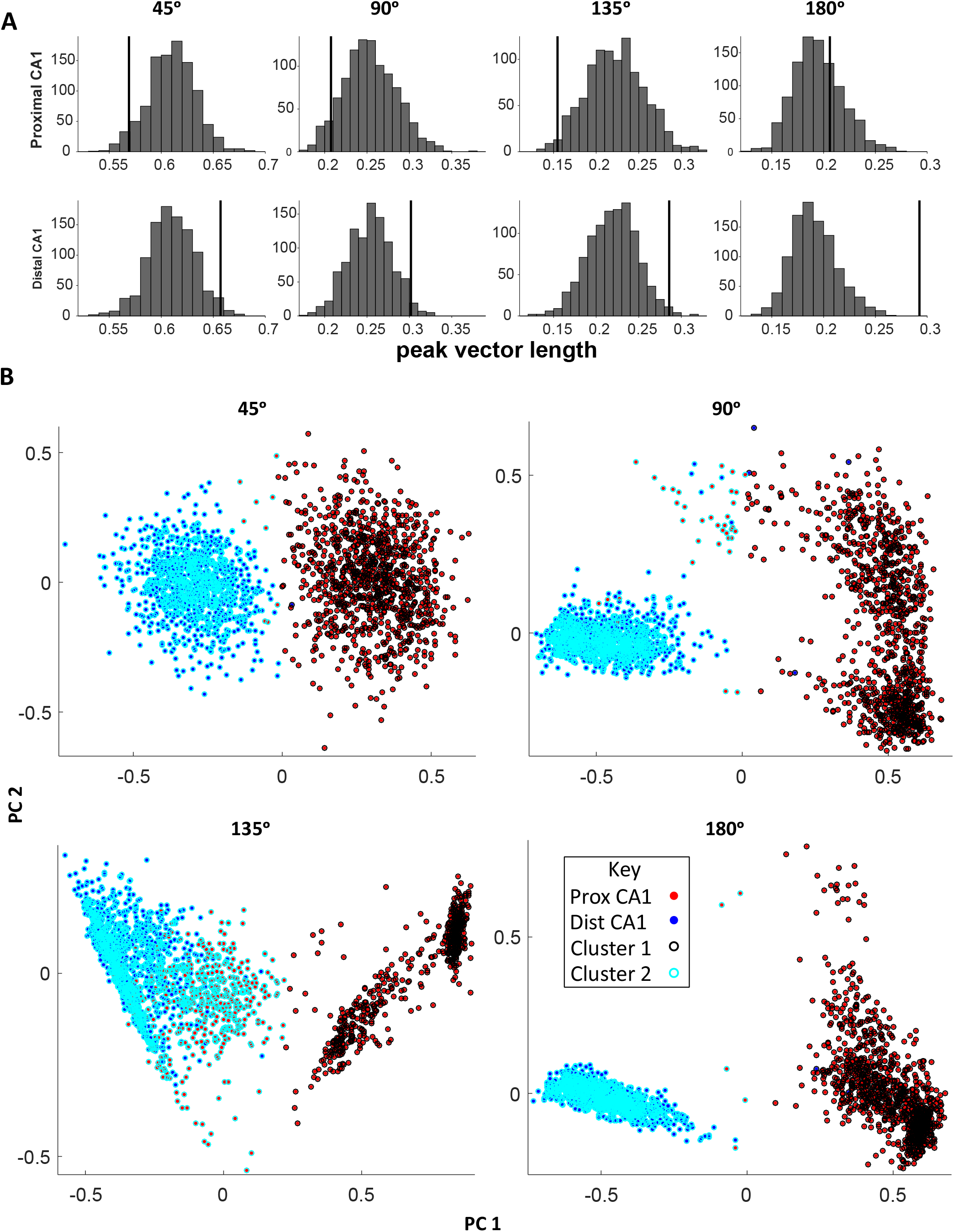
Shuffle and bootstrap analyses for PVCs. (A) Distributions of 1000 PVCs obtained by randomly reassigning neurons to pCA1 and dCA1 are shown in each histogram. Vertical lines mark the PVCs observed in real data. dCA1 has higher PVCs than the 95^th^ percentile of the shuffled distributions for all MIS angles, while pCA1 does not. (B) Bootstrapped distributions were created for pCA1 and dCA1 by resampling, with replacement, neurons in the two regions 1000 times. Number of samples in each iteration matched the number of neurons in the actual dataset. Peak vectors (x,y pairs), FWHM, and normalized bias were measured from the mean PVCs for all iterations. For each mismatch angle, PCA was run on this 4-dimensional space and the first two principal components were plotted to enable visualization of the bootstrapped distribution for pCA1 (red dots) and dCA1 (blue dots). Notice tighter clustering of dCA1 points compared to pCA1 points, which often form multiple clusters. k-means clustering (k = 2) was performed on the 4-dimensional data to classify the bootstrapped distributions into two clusters (black and cyan circles around the points). Almost all of the dCA1 points belong to one cluster while almost all of the pCA1 points belong to the other cluster for all mismatch angles other than 135° (see Supplementary Table 5 for the numbers of points from each region belonging to each cluster). At 135°, pCA1 points almost evenly divide into the two clusters.

To compare pCA1 and dCA1 PVCs explicitly, repeated sampling with replacement of neurons from each region was performed 1000 times to generate bootstrapped distributions for the two regions. The number of samples in each iteration matched the number of neurons in the real dataset for each subregion. These bootstraps allow us to estimate sample distribution of the PVC. For each of the iterations, PVC matrices were generated and mean PVC polar plots for each relative displacement were computed from those matrices. The following parameters were estimated from the polar plots to facilitate comparison between pCA1 and dCA1: Peak vectors (x,y pairs), FWHM, and normalized bias (see methods for definitions of and calculations of these parameters). PCA was performed on this 4-dimensional space for reducing dimensionality for display purposes. Figure 6B shows the projection of the bootstrapped data for the two regions on the first two principal components for the different mismatch angles. pCA1 bootstraps are shown in red and dCA1 bootstraps are shown in blue. Three clear patterns stand out in all four plots. First, the two distributions are well segregated along the two principal components, with the first principal component showing a clear segregation between the two at all mismatch angles other than 135°. This segregation indicates that pCA1 and dCA1 mean PVC distributions are different from one another. Second, the dCA1 bootstrap distribution is much more compact at all mismatch angles than the pCA1 bootstrap distribution. Third, the pCA1 bootstrap distribution often shows multiple clusters, while almost all dCA1 bootstraps stay together in a single compact cluster. In fact, the pCA1 bootstrap distribution for 135° mismatch shows 3 clusters with two clusters clearly segregated from the dCA1 cluster and the third cluster overlapping with the dCA1 cluster. The multiple modes in the pCA1 distributions indicate that the underlying population has multiple competing rotational propensities of comparable magnitudes in response to cue conflict. In contrast, compact unimodal clusters in the dCA1 bootstrapped distribution indicate that the dCA1 population responds more coherently to cue conflict. This observation corroborates the pattern of responses seen in the PVC matrices in Figure 5.

To test whether these distributions of pCA1 and dCA1 bootstraps are indeed different from one another in an unbiased manner, k-means clustering with k = 2 was employed. Figure 6B shows black and cyan circles surrounding points classified into two clusters. Supplementary Table 5 shows numbers of points from pCA1 and dCA1 that were included in the two clusters. At 135° mismatch, all the dCA1 points and a group of pCA1 points clustered together, while the remaining pCA1 points formed the other cluster. For all other mismatch angles, the clusters include more than 95% of the points from one region while including less than 5% of the points from the other region, as expected. These clustering patterns prove that pCA1 and dCA1 respond differentially to the double rotation cue manipulation, and that dCA1 population rotates more coherently than the pCA1 population.

## Discussion

### Dissociation of responses to double rotation manipulation between pCA1 and dCA1

EC and CA3 inputs to pCA1 and dCA1 respond differently to double rotation, leading to an expectation that the responses to this manipulation will differ along the transverse axis of CA1. pCA1 receives strong competing inputs from dCA3 and MEC during MIS sessions. dCA3 rotates coherently with local cues (Lee et al. 2015), while MEC rotates coherently with global cues (Neunuebel et al. 2013). In contrast, dCA1 does not receive a strong, coherently rotating signal from either CA3 or LEC. pCA3 responds incoherently to double rotation, with sporadic hotspots of high correlation in the PVC matrices of MIS angles > 45° rather than a band of high correlation following either the local or the global cues (Lee et al. 2015). The sporadic hotspots of high correlation are sometimes seen rotating with local cues while rotating with global cues at some other times. LEC neurons show a weak spatial selectivity on the circular track in STD sessions (Yoganarasimha et al. 2011). Neither the STD1 vs STD2 nor the STD1 vs MIS PVC matrices for LEC show a narrow band of high correlation, but polar plots of mean PVCs reveal a weak but statistically significant preference for rotating with local cues (Neunuebel et al. 2013). LEC encodes external items in egocentric coordinates (Wang et al. 2018), but in the present task, the allocentric and the egocentric reference frames are confounded, since the rat typically faces in a certain direction at each position on track. This translates into distances and directions to different cues being practically fixed at each position. Hence, we do not expect the response of the LEC neurons to double rotation manipulation to appear different between the two reference frames and expect the local cues to weakly dominate over global cues (Neunuebel et al. 2013) even in the egocentric reference frame.

These weakly local and incoherent inputs from LEC and pCA1 would predict either incoherent or weakly local response in dCA1. Surprisingly, dCA1 shows a more coherent rotation than pCA1 in this study, rotating with global cues at all MIS angles. Alternative sources of spatial information anchored to global cues need to be considered to explain these results. One possibility is that in absence of strong competing inputs, even sparse projections from MEC (Masurkar et al. 2017) might be sufficient to cause dCA1 neurons to rotate with global cues. In addition to EC inputs, CA1 receives other inputs which could potentially influence spatial representation in dCA1 in this task. Nucleus reuniens, which connects to the medial prefrontal cortex and the hippocampus (Dolleman-van der Weel et al. 2019; Dolleman-Van Der Weel and Witter 1996; Vertes et al. 2006), sends direct projections to CA1 and shows head direction cells (Jankowski et al. 2014), place cells, and border cells (Jankowski et al. 2015). While nucleus reuniens neurons have not been recorded in the double rotation task, head direction cells recorded in other thalamic nuclei (ADN, AVN, LDN, VAN and RT) rotate with the global cues in the double rotation task (Yoganarasimha et al. 2006). Head direction selectivity has been reported in CA1 (Leutgeb et al. 2000). Anecdotally, we as well as others (Peyrache and Buzsaki, https://www.youtube.com/watch?v&=2da6gzF9eo0&t=24m13s) have recorded putative axons with head direction tuning outside pyramidal cell layer in dCA1. Notwithstanding the source of these head direction projections, this anecdotal observation further supports the hypothesis that head direction inputs may be involved in coherent rotation of dCA1 with global cues. Subiculum, which receives direct projections from the head direction system as well as MEC (Ding 2013), sends feedback projections to CA1 (Xu et al. 2016), and rotates coherently with global cues in the double rotation task (Sharma et al. 2020). While these inputs from MEC, nucleus reuniens, and the subiculum to dCA1 may be less numerous than LEC and pCA3, they may be strong enough to drive dCA1 to rotate with global cues in absence of competing strong inputs rotating with local cues.

In contrast to dCA1, pCA1 gets strong, conflicting inputs from dCA3 (Lee et al. 2015) and MEC (Neunuebel et al. 2013) rotating with local and global cues respectively during double rotation. This incoherent response may be explained by differential strengths of CA3 and MEC inputs to individual pCA1 neurons driving the variety of responses observed there. Neurons with predominant inputs from either CA3 or MEC may follow that input, while those without a clearly dominant input from CA3 and MEC may give ambiguous or remapping responses.

### Spatial selectivity in pCA1 and dCA1

In this study, putative pyramidal cells in pCA1 and dCA1 show comparable spatial selectivity as estimated using spatial information score, number of place fields per cell, field size, and fraction of the track occupied by fields. This lack of difference in spatial selectivity along the transverse axis observed in STD as well as MIS sessions contradicts prior studies showing higher spatial selectivity in pCA1 compared to dCA1 (Henriksen et al. 2010; Ng et al. 2018; Oliva et al. 2016). The behavioral arenas used in the earlier studies had a single uniform texture, while the circular track in the present study had 4 easily discernible sections with distinct textures, visual appearances, and odors. Thus, the apparent elimination of differences in spatial selectivity between pCA1 and dCA1 could possibly be caused by richer sensory information available from the behavioral arena. This observation challenges the established notion that dCA1 is less spatial than pCA1. Instead, the functional difference between pCA1 and dCA1 appears to be caused by the kind of inputs used for creating spatial representations in the two areas. More coherent rotations in response to cue manipulations seen in dCA1 than pCA1 corroborate this claim of dCA1 being at least as spatial as pCA1.

LEC shows weak spatial selectivity on the circular track used here, similar to that in a square box in simple and complex environments (Hargreaves et al. 2005; Neunuebel et al. 2013; Yoganarasimha et al. 2011). Hence, spatial selectivity of the inputs from LEC on the circular track is insufficient to explain the higher spatial selectivity in dCA1 in the present study, and projections from MEC (Masurkar et al. 2017), nucleus reuniens (Dolleman-van der Weel et al. 2019; Dolleman-Van Der Weel and Witter 1996; Vertes et al. 2006), and the subiculum (Ding 2013; Xu et al. 2016) to dCA1 discussed earlier need to be considered. However, these inputs may not be sufficient on their own to explain why the spatial selectivity differences along the transverse axis of CA1 seen in the previous studies (Henriksen et al. 2010; Ng et al. 2018; Oliva et al. 2016) disappear in the present study since these inputs would have been available to dCA1 in the earlier experiments, too. It is possible that the availability of the on-track sensory cues increases spatial selectivity of the non-LEC inputs on the circular track compared to the other environments used in the earlier studies. Alternatively, increased complexity of the environment may increase spatial encoding demands on the hippocampal system, which may respond by recruiting dCA1 in a task dependent manner. This can be achieved by increasing responsiveness of dCA1 to pre-existing spatial inputs with increasing complexity. Further studies are required to test if one or both of these mechanisms contribute to increased spatial selectivity in dCA1, including the role of LEC as the possible modulator of contributions of non-LEC inputs to the CA1 spatial code (Lu et al. 2013). Thus, it is clear that the functional distinction between pCA1 and dCA1 is more complex than merely being more or less spatial and many more inputs beyond LEC and MEC may play a role in generating distinct spatial representations in pCA1 and dCA1.

### Reference frames in CA1

Since the discovery of place cells (O’Keefe 1976; O’Keefe and Dostrovsky 1971), spatial representation in the hippocampus is thought to be a unitary representation of absolute space (O’keefe and Nadel 1978). This unitary representation requires a single reference frame with a single origin with respect to which every point of space is encoded. However, it is possible that the hippocampal spatial representation might appear to be unitary, without there being an explicitly unitary representation of space. A number of independent reference frames/spatial representations may coexist, and may generally agree with one another, giving an appearance of a unitary code for space. Thus, different neurons firing at a specific location may be encoding that location as different distances and directions from different origins. Some experimental conditions demonstrate existence of such coexisting reference frames in CA1.

Representations of proximity to barriers (Rivard et al. 2004), goal location (Fyhn et al. 2002; Gothard et al. 1996b), distance and direction to goal (Gothard et al. 1996a; Sarel et al. 2017), distance and direction to landmarks (Deshmukh and Knierim 2013; McNaughton et al. 1995) may coexist with the classic place cells representing space in allocentric coordinates. These place fields in allocentric coordinates themselves may be formed by combining inputs from two or more boundary vector cells (Hartley et al. 2000; O’Keefe and Burgess 1996) or grid cells (de Almeida et al. 2009; Monaco et al. 2011; Savelli and Knierim 2010; Solstad et al. 2006), or a combination of sensory and self-generated inputs (Deshmukh and Knierim 2013). Thus, even the classic place cells may be encoding space in different reference frames.

Creating an explicit conflict between reference frames reveals their influence on hippocampal representation of space (Gothard et al. 1996a, 1996b; Knierim 2002; Lee et al. 2004; Zinyuk et al. 2000). In the double rotation paradigm used in this paper, some regions involved in spatial encoding show coherent rotations while the others show weak to no coherence in rotation (Lee et al. 2015, 2004; Neunuebel et al. 2013; Neunuebel and Knierim 2014; Sharma et al. 2020). The dissociation of responses to the double rotation manipulation along the proximodistal axis of CA1 shown here demonstrates that the non-unitary representation of space in the hippocampal output (via CA1) reflects strong conflict apparent in its inputs (in pCA1) or lack thereof (in dCA1). Regions of the brain differentially targeted (directly or indirectly) by pCA1 and dCA1 (Naber et al. 2001; Witter and Amaral 2004), thus, receive spatial inputs that have differential influence of different spatial reference frames. Further studies are required to elaborate the role of this difference of spatial inputs in functioning of these target regions.

## Acknowledgements

I thank Jeremy Johnson, Geeta Rao, Vyash Puliyadi, Amanda Smolinsky, Lou Blanpain, and Kimberley Christian for their support in data collection and James J Knierim for scientific discussions. This work was supported by Wellcome Trust/DBT India Alliance Grant IA/S/13/2/501024. Data collection was supported by R01 NS039456 (J. Knierim, PI).

