## Supplementary Information for "Distal CA1 maintains a more coherent spatial representation than proximal CA1 when local and global cues conflict"

### Supplementary Figure 1

45° Mismatch and preceding STD

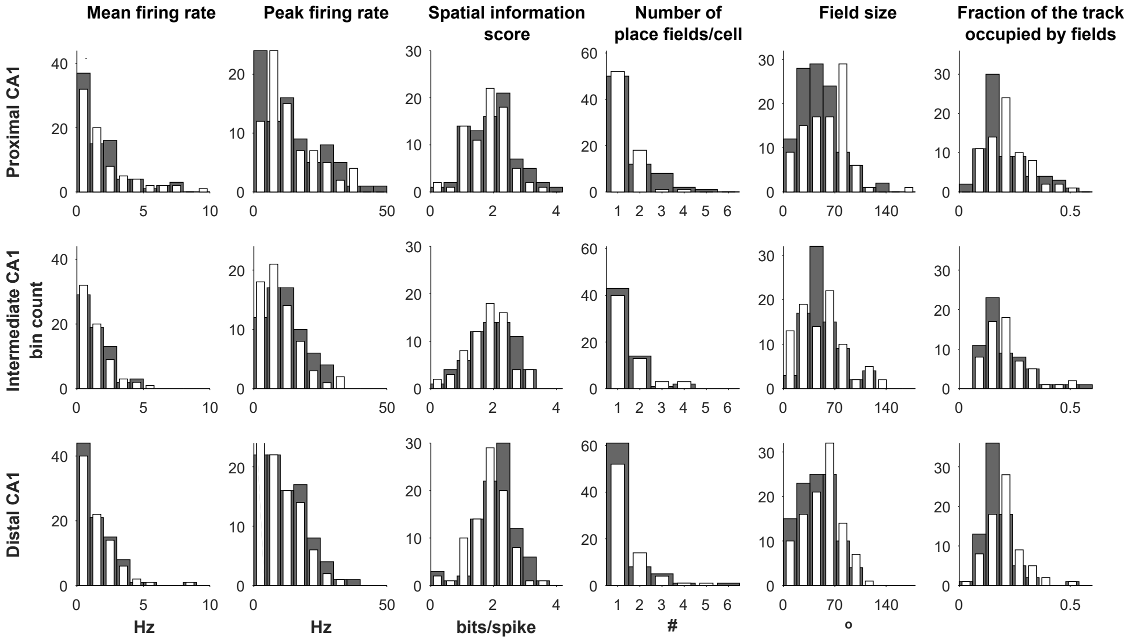


90°Mismatch and preceding STD

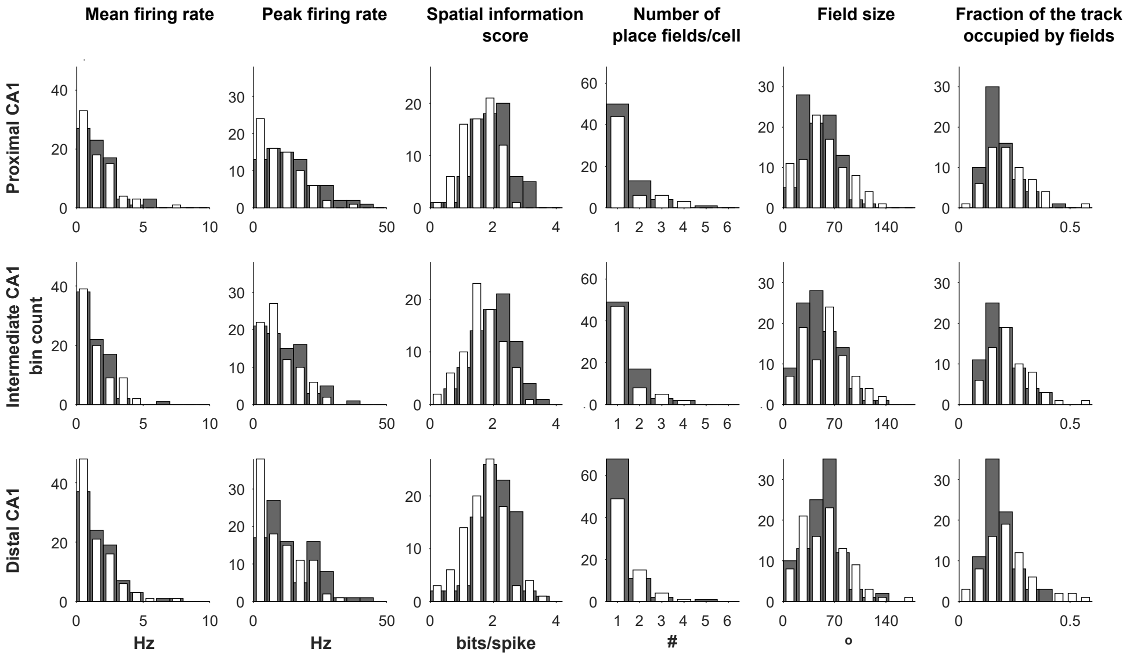


**Continued on the next page**

### Supplementary Figure 1 (cont.)

135° Mismatch and preceding STD

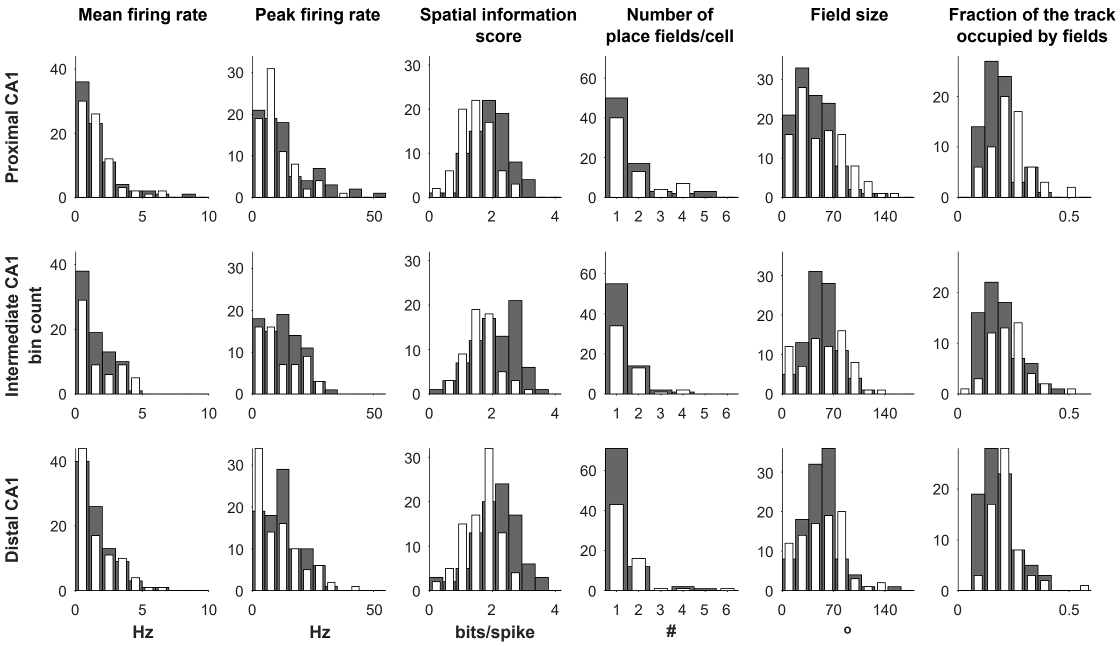


180°Mismatch and preceding STD

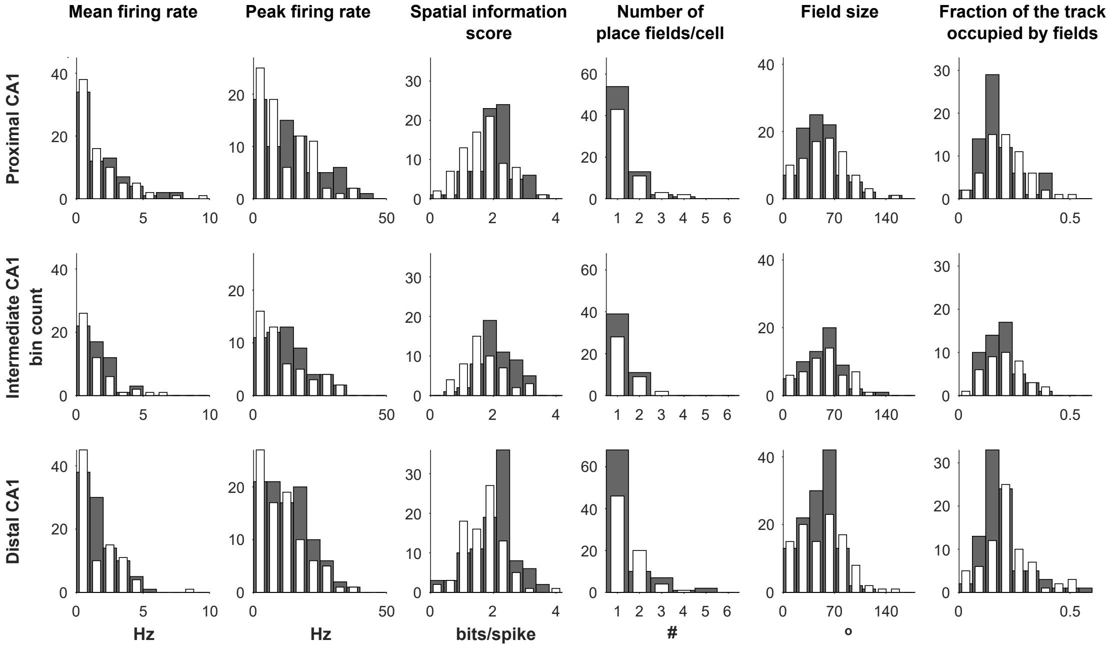


**Supplementary Figure 1. Modulation of properties of putative pyramidal cells along the transverse axis of CA1 by double rotation manipulation.** Mean firing rates, peak firing rates, spatial information scores, number of place fields per cell, place field sizes, and fraction of the track occupied by place fields in the proximal, intermediate, and distal CA1 during the various MIS sessions (white bars) and the STD sessions preceding them (gray bars) show that these parameters remain comparable across the transverse axis of CA1. A trend of reduced spatial selectivity from STD to MIS sessions can be seen in all three subregions.

**Tables**

| Statistical test | Quantities compared | p | Z | Wilcoxon ranksum | χ2 | Prox CA1 n | Prox CA1 median | Dist CA1 n | Dist CA1 median |
|---|---|---|---|---|---|---|---|---|---|
| Wilcoxon rank sum | Mean firing rate | 0.37 | 0.89 | 27079 | | 156 neurons | 1.24 Hz | 180 neurons | 1.29 Hz |
| Wilcoxon rank sum | Peak firing rate | 0.34 | 0.95 | 27128 | | 156 neurons | 11.97 Hz | 180 neurons | 11.52 Hz |
| Wilcoxon rank sum | Spatial information score | 0.15 | -1.45 | 24997 | | 156 neurons | 1.92 bits/spike | 180 neurons | 2.11 bits/spike |
| Wilcoxon rank sum | Number of place fields/cell | 0.28 | 1.07 | 19968 | | 139 place cells | 1 place field/cell | 140 place cells | 1 place field/cell |
| Wilcoxon rank sum | Field size | 0.13 | -1.52 | 30915 | | 181 place fields | 44° | 176 place fields | 47° |
| Wilcoxon rank sum | Fraction of area occupied by place fields | 0.78 | -0.28 | 19272 | | 139 place cells | 0.17 | 140 place cells | 0.17 |
| χ2 | Fraction of place cells with 1 place field/cell | 0.36 | | | 0.85 | 107 place cells with 1 field out of 139 place cells | | 115 place cells with 1 field out of 140 place cells | |

**Supplementary Table 1, related to Figure 2**. **Statistical comparison of properties of proximal and distal CA1 neurons in the first standard session of the day.**

| Session | Statistical test | Quantities compared | Regions & sessions compared | p | Z | Wilcoxon ranksum | χ2 | Prox CA1 n, median | Dist CA1 n, median |
|---|---|---|---|---|---|---|---|---|---|
| 45° MIS & STD preceding | Wilcoxon rank sum | Mean firing rate | Proximal vs Distal (MIS vs MIS) | 0.0768 | 1.7737 | 6723 | | 76, 1.26 Hz | 86, 1.13 Hz |
| | Wilcoxon rank sum | Mean firing rate | Proximal STD vs MIS | 0.3501 | -0.9344 | 6250 | | MIS = 76, 1.26 Hz STD = 82, 1.24 Hz | |
| | Wilcoxon rank sum | Mean firing rate | Distal STD vs MIS | 0.7635 | -0.3008 | 7996 | | | MIS = 86, 1.13 Hz STD = 91, 1.08 Hz |
| | Wilcoxon rank sum | Peak firing rate | Proximal vs Distal (MIS vs MIS) | 0.0767 | 1.7704 | 6722 | | 76, 10.78 Hz | 86, 9.62 Hz |
| | Wilcoxon rank sum | Peak firing rate | Proximal STD vs MIS | 0.6397 | -0.4681 | 6384 | | MIS = 76, 10.78 Hz STD = 82, 10.99 Hz | |
| | Wilcoxon rank sum | Peak firing rate | Distal STD vs MIS | 0.4021 | 0.8379 | 8385 | | | MIS = 86, 9.62 Hz STD = 91, 10.17 Hz |
| | Wilcoxon rank sum | Spatial information score | Proximal vs Distal (MIS vs MIS) | 0.4862 | -0.6964 | 5986 | | 76, 1.85 b/s | 86, 1.88 b/s |
| | Wilcoxon rank sum | Spatial information score | Proximal STD vs MIS | 0.2647 | 1.1153 | 6840 | | MIS = 76, 1.85 b/s STD = 82, 2 b/s | |
| | Wilcoxon rank sum | Spatial information score | Distal STD vs MIS | 0.007 | 2.6958 | 9018 | | | MIS = 86, 1.88 b/s STD = 91, 2.14 b/s |
| | Wilcoxon rank sum | Number of place fields/cell | Proximal vs Distal (MIS vs MIS) | 0.8422 | -0.1991 | 5180.5 | | 72, 1 | 72, 1 |
| | Wilcoxon rank sum | Number of place fields/cell | Proximal STD vs MIS | 0.3681 | 0.9001 | 5512 | | MIS = 72, 1 STD = 73, 1 | |
| | Wilcoxon rank sum | Number of place fields/cell | Distal STD vs MIS | 0.3137 | -1.0074 | 5466.5 | | | MIS = 72, 1 STD = 76, 1 |
| | Wilcoxon rank sum | Field size | Proximal vs Distal (MIS vs MIS) | 0.1367 | 1.4881 | 9948.5 | | 95, 52° | 101, 49° |
| | Wilcoxon rank sum | Field size | Proximal STD vs MIS | 0.0024 | -3.0355 | 10194 | | MIS = 95, 52° STD = 111, 40° | |
| | Wilcoxon rank sum | Field size | Distal STD vs MIS | 0.0244 | -2.2513 | 9461.5 | | | MIS = 101, 49° STD = 102, 38° |
| | Wilcoxon rank sum | Fraction of area occupied by place fields | Proximal vs Distal (MIS vs MIS) | 0.653 | 0.4496 | 5333 | | 72, 0.21 | 72, 0.19 |
| | Wilcoxon rank sum | Fraction of area occupied by place fields | Proximal STD vs MIS | 0.2531 | -1.143 | 5039.5 | | MIS = 72, 0.21 STD = 73, 0.17 | |
| | Wilcoxon rank sum | Fraction of area occupied by place fields | Distal STD vs MIS | 5.04E-04 | -3.4785 | 4755 | | | MIS = 72, 0.19 STD = 76, 0.16 |
| | χ2 | Fraction of place cells | Proximal vs Distal (MIS vs MIS) | 0.8524 | | | 0.0346 | 45/72 | 42/72 |

| | | | | | | | | | |
|---|---|---|---|---|---|---|---|---|---|
| | | with 1 place field/cell | | | | | | | |
| | χ2 | Fraction of place cells with 1 place field/cell | Proximal STD vs MIS | 0.7568 | | | 0.0959 | MIS = 45/72 STD = 45/73 | |
| | χ2 | Fraction of place cells with 1 place field/cell | Distal STD vs MIS | 0.3385 | | | 0.9161 | | MIS = 42/72 STD = 50/76 |
| **Session** | **Statistical test** | **Quantities compared** | **Regions & sessions compared** | **p** | **Z** | **Wilcoxon ranksum** | **χ2** | **# of samples in Prox CA1** | **# of samples in Dist CA1** |
| 90° MIS & STD preceding | Wilcoxon rank sum | Mean firing rate | Proximal vs Distal (MIS vs MIS) | 0.2014 | 1.2776 | 6734 | | 74, 1.13 Hz | 96, 1 Hz |
| | Wilcoxon rank sum | Mean firing rate | Proximal STD vs MIS | 0.4351 | 0.7804 | 5717 | | MIS=74, 1.13 Hz STD = 74, 1.41 Hz | |
| | Wilcoxon rank sum | Mean firing rate | Distal STD vs MIS | 0.0622 | 1.8648 | 9390 | | | MIS = 96, 1 Hz STD = 92, 1.27 Hz |
| | Wilcoxon rank sum | Peak firing rate | Proximal vs Distal (MIS vs MIS) | 0.4274 | 0.7936 | 6580 | | 74, 8.13 Hz | 96, 8.3 Hz |
| | Wilcoxon rank sum | Peak firing rate | Proximal STD vs MIS | 0.0298 | 2.1725 | 6080 | | MIS=74, 8.13 Hz STD = 74, 11.56 Hz | |
| | Wilcoxon rank sum | Peak firing rate | Distal STD vs MIS | 0.0047 | 2.8273 | 9749 | | | MIS = 96, 8.3 Hz STD = 92, 10.58 Hz |
| | Wilcoxon rank sum | Spatial information score | Proximal vs Distal (MIS vs MIS) | 0.11 | -1.5982 | 5818 | | 74, 1.8 b/s | 96, 1.61 b/s |
| | Wilcoxon rank sum | Spatial information score | Proximal STD vs MIS | 1.98E-04 | 3.7219 | 6484 | | MIS=74, 1.8 b/s STD = 74, 2.02 b/s | |
| | Wilcoxon rank sum | Spatial information score | Distal STD vs MIS | 0.0011 | 3.2563 | 9909 | | | MIS = 96, 1.61 b/s STD = 92, 2.07 b/s |
| | Wilcoxon rank sum | Number of place fields/cell | Proximal vs Distal (MIS vs MIS) | 0.9197 | -0.1009 | 3788.5 | | 59, 1 | 69, 1 |
| | Wilcoxon rank sum | Number of place fields/cell | Proximal STD vs MIS | 0.8825 | -0.1478 | 4328 | | MIS = 59, 1 STD = 68, 1 | |
| | Wilcoxon rank sum | Number of place fields/cell | Distal STD vs MIS | 0.081 | -1.7451 | 5891.5 | | | MIS = 69, 1 STD = 82, 1 |
| | Wilcoxon rank sum | Field size | Proximal vs Distal (MIS vs MIS) | 0.6311 | -0.4802 | 7656.5 | | 86, 45° | 95, 53° |
| | Wilcoxon rank sum | Field size | Proximal STD vs MIS | 0.2203 | -1.2258 | 7945 | | MIS = 86, 45° STD = 93, 41° | |
| | Wilcoxon rank sum | Field size | Distal STD vs MIS | 0.5403 | -0.6124 | 9705 | | | MIS = 95, 53° STD = 101, 48° |
| | Wilcoxon rank sum | Fraction of area occupied by place fields | Proximal vs Distal (MIS vs MIS) | 0.7128 | 0.3681 | 3883 | | 59, 0.21 | 69, 0.2 |

| | | | | | | | | | |
|---|---|---|---|---|---|---|---|---|---|
| | Wilcoxon rank sum | Fraction of area occupied by place fields | Proximal STD vs MIS | 0.0193 | -2.34 | 3867.5 | | MIS = 59, 0.21 STD = 68, 0.17 | |
| | Wilcoxon rank sum | Fraction of area occupied by place fields | Distal STD vs MIS | 0.056 | -1.911 | 5720 | | | MIS = 69, 0.2 STD = 82, 0.17 |
| | χ2 | Fraction of place cells with 1 place field/cell | Proximal vs Distal (MIS vs MIS) | 0.8012 | | | 0.0634 | 35/59 | 42/69 |
| | χ2 | Fraction of place cells with 1 place field/cell | Proximal STD vs MIS | 0.9452 | | | 0.0047 | MIS = 35/59, STD = 42/68 | |
| | χ2 | Fraction of place cells with 1 place field/cell | Distal STD vs MIS | 0.1211 | | | 2.4031 | | Mis = 42/69 STD = 58/82 |
| **Session** | **Statistical test** | **Quantities compared** | **Regions & sessions compared** | **p** | **Z** | **Wilcoxon ranksum** | **χ2** | **# of samples in Prox CA1** | **# of samples in Dist CA1** |
| 135° MIS & STD preceding | Wilcoxon rank sum | Mean firing rate | Proximal vs Distal (MIS vs MIS) | 0.5473 | 0.6018 | 6453 | | 76, 1.15 Hz | 88, 1.01 Hz |
| | Wilcoxon rank sum | Mean firing rate | Proximal STD vs MIS | 0.8274 | -0.2181 | 6218 | | MIS = 76, 1.15 Hz STD = 80, 1.16 Hz | |
| | Wilcoxon rank sum | Mean firing rate | Distal STD vs MIS | 0.4161 | 0.8132 | 8750 | | | MIS = 88, 1.01 Hz STD = 93, 1.19 Hz |
| | Wilcoxon rank sum | Peak firing rate | Proximal vs Distal (MIS vs MIS) | 0.9882 | -0.0148 | 6265 | | 76, 7.82 Hz | 88, 8.54 Hz |
| | Wilcoxon rank sum | Peak firing rate | Proximal STD vs MIS | 0.1134 | 1.5831 | 6727 | | MIS = 76, 7.82 Hz STD = 80, 10.16 Hz | |
| | Wilcoxon rank sum | Peak firing rate | Distal STD vs MIS | 0.0628 | 1.8606 | 9119 | | | MIS = 88, 8.54 Hz STD = 93, 12.29 Hz |
| | Wilcoxon rank sum | Spatial information score | Proximal vs Distal (MIS vs MIS) | 0.0125 | -2.4979 | 5512 | | 76, 1.52 b/s | 88, 1.77 b/s |
| | Wilcoxon rank sum | Spatial information score | Proximal STD vs MIS | 7.23E-06 | 4.4869 | 7546 | | MIS = 76, 1.52 b/s STD = 80, 1.95 b/s | |
| | Wilcoxon rank sum | Spatial information score | Distal STD vs MIS | 3.22E-06 | 4.6564 | 10104 | | | MIS = 88, 1.77 b/s STD = 93, 2.14 b/s |
| | Wilcoxon rank sum | Number of place fields/cell | Proximal vs Distal (MIS vs MIS) | 0.244 | 1.1651 | 4264.5 | | 64, 1 | 62, 1 |
| | Wilcoxon rank sum | Number of place fields/cell | Proximal STD vs MIS | 0.5136 | -0.6532 | 5118.5 | | MIS = 64, 1 STD = 75, 1 | |
| | Wilcoxon rank sum | Number of place fields/cell | Distal STD vs MIS | 0.0671 | -1.831 | 6061.5 | | | MIS = 62, 1 STD = 86, 1 |
| | Wilcoxon rank sum | Field size | Proximal vs Distal (MIS vs MIS) | 0.4952 | -0.682 | 10069 | | 106, 37° | 88, 51° |

| Session | Statistical test | Quantities compared | Regions & sessions compared | p | Z | Wilcoxon ranksum | χ2 | # of samples in Prox CA1 | # of samples in Dist CA1 |
|---|---|---|---|---|---|---|---|---|---|
| | Wilcoxon rank sum | Field size | Proximal STD vs MIS | 0.0566 | -1.9061 | 12023 | | MIS = 106, 37° STD = 116, 35° | |
| | Wilcoxon rank sum | Field size | Distal STD vs MIS | 0.4998 | -0.6748 | 10371 | | | MIS = 88, 51° STD = 108, 43° |
| | Wilcoxon rank sum | Fraction of area occupied by place fields | Proximal vs Distal (MIS vs MIS) | 0.0358 | 2.0988 | 4494.5 | | 64, 0.23 | 62, 0.2 |
| | Wilcoxon rank sum | Fraction of area occupied by place fields | Proximal STD vs MIS | 3.51E-05 | -4.1377 | 4270.5 | | MIS = 64, 0.23 STD = 75, 0.17 | |
| | Wilcoxon rank sum | Fraction of area occupied by place fields | Distal STD vs MIS | 0.0018 | -3.1156 | 5605 | | | MIS = 62, 0.2 STD = 86, 0.17 |
| | χ2 | Fraction of place cells with 1 place field/cell | Proximal vs Distal (MIS vs MIS) | 0.533 | | | 0.3886 | 34/64 | 37/62 |
| | χ2 | Fraction of place cells with 1 place field/cell | Proximal STD vs MIS | 0.7381 | | | 0.1118 | MIS = 34/64 STD = 45/75 | |
| | χ2 | Fraction of place cells with 1 place field/cell | Distal STD vs MIS | 0.0918 | | | 2.8423 | | MIS = 37/62 STD = 60/85 |
| 180° MIS & STD preceding | Wilcoxon rank sum | Mean firing rate | Proximal vs Distal (MIS vs MIS) | 0.7859 | 0.2716 | 6518 | | 78, 1.1 Hz | 86, 0.95 Hz |
| | Wilcoxon rank sum | Mean firing rate | Proximal STD vs MIS | 0.5778 | 0.5566 | 5928 | | MIS =78, 1.1 Hz STD = 75 1.3 Hz | |
| | Wilcoxon rank sum | Mean firing rate | Distal STD vs MIS | 0.5876 | 0.5424 | 9261 | | | MIS = 86, 0.95 Hz STD = 98 1.16 Hz |
| | Wilcoxon rank sum | Peak firing rate | Proximal vs Distal (MIS vs MIS) | 0.7283 | 0.3474 | 6541 | | 78, 7.31 Hz | 86, 9.8 Hz |
| | Wilcoxon rank sum | Peak firing rate | Proximal STD vs MIS | 0.0846 | 1.7245 | 6248 | | MIS =78, , 7.31 Hz STD = 75, 12.02 Hz | |
| | Wilcoxon rank sum | Peak firing rate | Distal STD vs MIS | 0.0669 | 1.8324 | 9726 | | | MIS = 86, , 9.8 Hz STD = 98, 11.91 Hz |
| | Wilcoxon rank sum | Spatial information score | Proximal vs Distal (MIS vs MIS) | 0.8939 | -0.1334 | 6394 | | 78, 1.68 b/s | 86, 1.71 b/s |
| | Wilcoxon rank sum | Spatial information score | Proximal STD vs MIS | 9.63E-05 | 3.8997 | 6844 | | MIS =78, 1.68 b/s STD = 75, 2.09 b/s | |
| | Wilcoxon rank sum | Spatial information score | Distal STD vs MIS | 3.22E-04 | 3.5968 | 10362 | | | MIS = 86, 1.71 b/s STD = 98, 2.11 b/s |
| | Wilcoxon rank sum | Number of place fields/cell | Proximal vs Distal (MIS vs MIS) | 0.4091 | -0.8255 | 3720 | | 59, 1 | 71, 1 |

| | | | | | | | | | |
|---|---|---|---|---|---|---|---|---|---|
| | Wilcoxon rank sum | Number of place fields/cell | Proximal STD vs MIS | 0.5143 | -0.6522 | 4445.5 | | MIS = 59, 1<br>STD = 70, 1 | |
| | Wilcoxon rank sum | Number of place fields/cell | Distal STD vs MIS | 0.1727 | -1.3636 | 6728.5 | | | MIS = 71, 1<br>STD = 88, 1 |
| | Wilcoxon rank sum | Field size | Proximal vs Distal (MIS vs MIS) | 0.5493 | 0.5988 | 7800.5 | | 82, 43° | 102, 42° |
| | Wilcoxon rank sum | Field size | Proximal STD vs MIS | 0.2348 | -1.1881 | 7397 | | MIS = 82, 43°<br>STD = 90, 44° | |
| | Wilcoxon rank sum | Field size | Distal STD vs MIS | 0.4331 | -0.784 | 13518 | | | MIS = 102, 42° STD = 123, 45° |
| | Wilcoxon rank sum | Fraction of area occupied by place fields | Proximal vs Distal (MIS vs MIS) | 0.8644 | 0.1707 | 3901.5 | | 59, 0.2 | 71, 0.19 |
| | Wilcoxon rank sum | Fraction of area occupied by place fields | Proximal STD vs MIS | 0.0096 | -2.5889 | 4002 | | MIS = 59, 0.2<br>STD = 70, 0.16 | |
| | Wilcoxon rank sum | Fraction of area occupied by place fields | Distal STD vs MIS | 0.0421 | -2.0324 | 6453 | | | MIS = 71, 0.19 STD = 88, 0.17 |
| | χ2 | Fraction of place cells with 1 place field/cell | Proximal vs Distal (MIS vs MIS) | 0.4243 | | | 0.6385 | 32/59 | 37/71 |
| | χ2 | Fraction of place cells with 1 place field/cell | Proximal STD vs MIS | 0.7236 | | | 0.1251 | MIS = 32/59<br>STD = 49/70 | |
| | χ2 | Fraction of place cells with 1 place field/cell | Distal STD vs MIS | 0.1187 | | | 2.4342 | | Mis = 37/71<br>STD = 58/88 |
| **Session** | **Statistical test** | **Quantities compared** | **Regions & sessions compared** | **p** | **Z** | **Wilcoxon ranksum** | **χ2** | **# of samples in Prox CA1** | **# of samples in Dist CA1** |

**Supplementary Table 2. Statistical comparison of properties of proximal and distal CA1 neurons in MIS sessions with each other and with preceding STD sessions**. Notice that the only persistent trend is that of reduced spatial selectivity in both proximal and distal CA1 MIS sessions compared to preceding STD sessions. b/s stands for bits/spike.

| Mismatch angle | Prox CA1 | | | Int CA1 | | | Dist CA1 | | |
|---|---|---|---|---|---|---|---|---|---|
| | U | Critical value of U for p < 0.001 | n | U | Critical value of U for p < 0.001 | n | U | Critical value of U for p < 0.001 | n |
| 45° | 20333.32 | 163.60 | 58 | 20298.83 | 170.54 | 49 | 20360.76 | 163.60 | 57 |
| 90° | 19622.83 | 172.58 | 41 | 19918.09 | 172.58 | 41 | 19408.52 | 170.54 | 49 |
| 135° | 20010.00 | 170.54 | 48 | 19098.39 | 172.58 | 41 | 18588.52 | 172.58 | 44 |
| 180° | 17530.99 | 177.88 | 35 | 18459.68 | 184.05 | 27 | 16786.14 | 163.60 | 55 |

**Supplementary Table 3. Rao's spacing test statistics (Rao 1976; Russell and Levitin 1995) for single unit rotation data shown in Figure 4.** All tests were significant with $p < 0.001$.

| Mis ang | Prox CA1 | | | | | Int CA1 | | | | | Dist CA1 | | | | |
|---|---|---|---|---|---|---|---|---|---|---|---|---|---|---|---|
| | MVA | MVL | p | z | n | MVA | MVL | p | z | n | MVA | MVL | p | z | n |
| 45° | -7° | 0.83 | $< 10^{-5}$ | 40.2 | 58 | 2° | 0.82 | $< 10^{-5}$ | 33.5 | 49 | -5° | 0.91 | $< 10^{-5}$ | 46.9 | 57 |
| 90° | 35° | 0.41 | 0.0009 | 6.8 | 41 | -13° | 0.67 | $< 10^{-5}$ | 18.3 | 41 | -14° | 0.56 | $< 10^{-5}$ | 15.3 | 49 |
| 135° | 16° | 0.25 | 0.0486 | 3.0 | 48 | -1° | 0.22 | 0.1331 | 2.1 | 41 | -45° | 0.29 | 0.0268 | 3.6 | 44 |
| 180° | 107° | 0.39 | 0.0131 | 4.3 | 35 | -80° | 0.15 | 0.5674 | 0.6 | 27 | -84° | 0.15 | 0.3085 | 1.2 | 55 |

**Supplementary Table 4. Rayleigh test for uniformity for single unit rotation data shown in Figure 4.** MVA and MVL denote mean vector angles and lengths, z denotes Rayleigh's z. Negative MVAs denote CW rotation in the direction of the global cues, while positive MVAs denote CCW rotation in the direction of the local cues.

| Mismatch angle | Region | Number of points in cluster 1 | Number of points in cluster 2 |
|---|---|---|---|
| 45 ° | Prox CA1 | 991 | 9 |
| | Dist CA1 | 1 | 999 |
| 90 ° | Prox CA1 | 963 | 37 |
| | Dist CA1 | 4 | 996 |
| 135 ° | Prox CA1 | 534 | 466 |
| | Dist CA1 | 0 | 1000 |
| 180 ° | Prox CA1 | 995 | 5 |
| | Dist CA1 | 2 | 998 |

**Supplementary Table 5. Number of points from proximal and distal CA1 assigned to clusters 1 and 2 by k-means clustering (k = 2).** For ease of visualization, after running the k-means clustering algorithm, the cluster with higher number of points from proximal CA1 was named cluster 1, while the other cluster with higher number of points from distal CA1 was named cluster 2.
